# Bacterial genes outnumber archaeal genes in eukaryotic genomes

**DOI:** 10.1101/779579

**Authors:** Julia Brückner, William F. Martin

## Abstract

The origin of eukaryotes is one of evolution’s most important transitions, yet it is still poorly understood. Evidence for how it occurred should be preserved in eukaryotic genomes. Based on phylogenetic trees from ribosomal RNA and ribosomal proteins, eukaryotes are typically depicted as branching together with or within archaea. This ribosomal affiliation is widely interpreted as evidence for an archaeal origin of eukaryotes. However, the extent to which the archaeal ancestry of genes for the cytosolic ribosomes of eukaryotic cells is representative for the rest of the eukaryotic genome is unknown. Here we have clustered 19,050,992 protein sequences from 5,443 bacteria and 212 archaea with 3,420,731 protein sequences from 150 eukaryotes spanning six eukaryotic supergroups to identify genes that link eukaryotes exclusively to bacteria and archaea respectively. By downsampling the bacterial sample we obtain estimates for the bacterial and archaeal proportions of genes among 150 eukaryotic genomes. Eukaryotic genomes possess a bacterial majority of genes. On average, eukaryotic genes are 56% bacterial in origin. The majority drops to 53% in eukaryotes that never possessed plastids, and increases to 61% in photosynthetic eukaryotic lineages, where the cyanobacterial ancestor of plastids contributed additional genes to the eukaryotic genome, reaching 67% in higher plants. Intracellular parasites, which undergo reductive evolution in adaptation to the nutrient rich environment of the cells that they infect, relinquish bacterial genes for metabolic processes. In the current sample, this process of adaptive gene loss is most pronounced in the human parasite Encephalitozoon intestinalis with 86% archaeal and 14% bacterial derived genes. The most bacterial eukaryote genome sampled is rice, with 67% bacterial and 33% archaeal genes. The functional dichotomy, initially described for yeast, of archaeal genes being involved in genetic information processing and bacterial genes being involved in metabolic processes is conserved across all eukaryotic supergroups.

## Introduction

Biologists recognize three kinds of cells in nature: bacteria, archaea and eukaryotes. The bacteria and archaea are prokaryotic in organization, having generally small cells on the order of 0.5–5 microns in size and ribosomes that translate nascent mRNA molecules as they are synthesized on DNA (cotranscriptional translation) [1]. Eukaryotic cells are generally much larger in size, more complex in organization and have larger genomes possessing introns that are removed (spliced) from the mRNA on spliceosomes [2]. Eukaryotic cells always harbor a system of internal membranes [3,4] that form the endoplasmic reticulum and the cell nucleus, where splicing takes place [5]. Furthermore, eukaryotes typically possess double membrane bounded bioenergetic organelles, mitochondria, which were present in the eukaryote common ancestor (LECA) [6,7], but have undergone severe reduction in some lineages [8,9]. In terms of timing during Earth history, it is generally agreed that the first forms of life on Earth were prokaryotes, with isotopic evidence for the existence of bacterial and archaeal metabolic processes tracing back to rocks 3.5 billion years of age [10,11] or older [12]. The microfossil record indicates that eukaryotes arose later, about 1.4 to 1.6 billion years ago [13], hence that eukaryotes arose from prokaryotes. Though eukaryotes are younger than prokaryotes, the nature of their phylogenetic relationship(s) to bacteria and archaea remains debated because of differing views about the evolutionary origin of eukaryotic cells.

In the traditional three domain tree of life, eukaryotes are seen as a sister group to archaea [14–16] (Fig. 1a). In newer two-domain trees, eukaryotes are viewed as branching from within the archaea [17,18] (Fig 1b). In both the two domain and the three domain hypotheses, this is often seen as evidence for “an archaeal origin” of eukaryotes [17,18] (Fig. 1a,b). Germane to an archaeal origin is the view that eukaryotes are archaea that became more complex by gradualist evolutionary processes such as point mutation and gene duplication [19,20]. Countering that view are two sets of observations relating to symbiogenesis (origin through symbiosis) for eukaryotes (Fig. 1c,d). First, the archaea that branch closest to eukaryotes in the most recent phylogenies are very small in size (0.5 μm), they lack any semblance of eukaryote-like cellular complexity, and they live in obligate association with bacteria [21], clearly implicating symbiosis [21] rather than point mutation as the driving force at the origin of the eukaryotic clade (Fig. 1c). Second, and with a longer history in the literature, are the findings that mitochondria trace to the eukaryote common ancestor [22,23] and that many genes in eukaryote genomes trace to gene transfers from endosymbiotic organelles [24,25]. A symbiogenic origin of eukaryotes would run counter to one of the key goals of phylogenetics, namely to place eukaryotes in a natural system of phylogenetic classification where all groups are named according to their position in a bifurcating tree. If eukaryotes arose via symbiosis of an archaeon (the host) and a bacterium (the mitochondrion), then eukaryotes would reside simultaneously on both the archaeal and the bacterial branches in phylogenetic schemes [26,27], whereby plants and algae that stem from secondary symbioses [28] would reside on recurrently anastomosing branches as in Fig. 1d.

**Figure 1.**
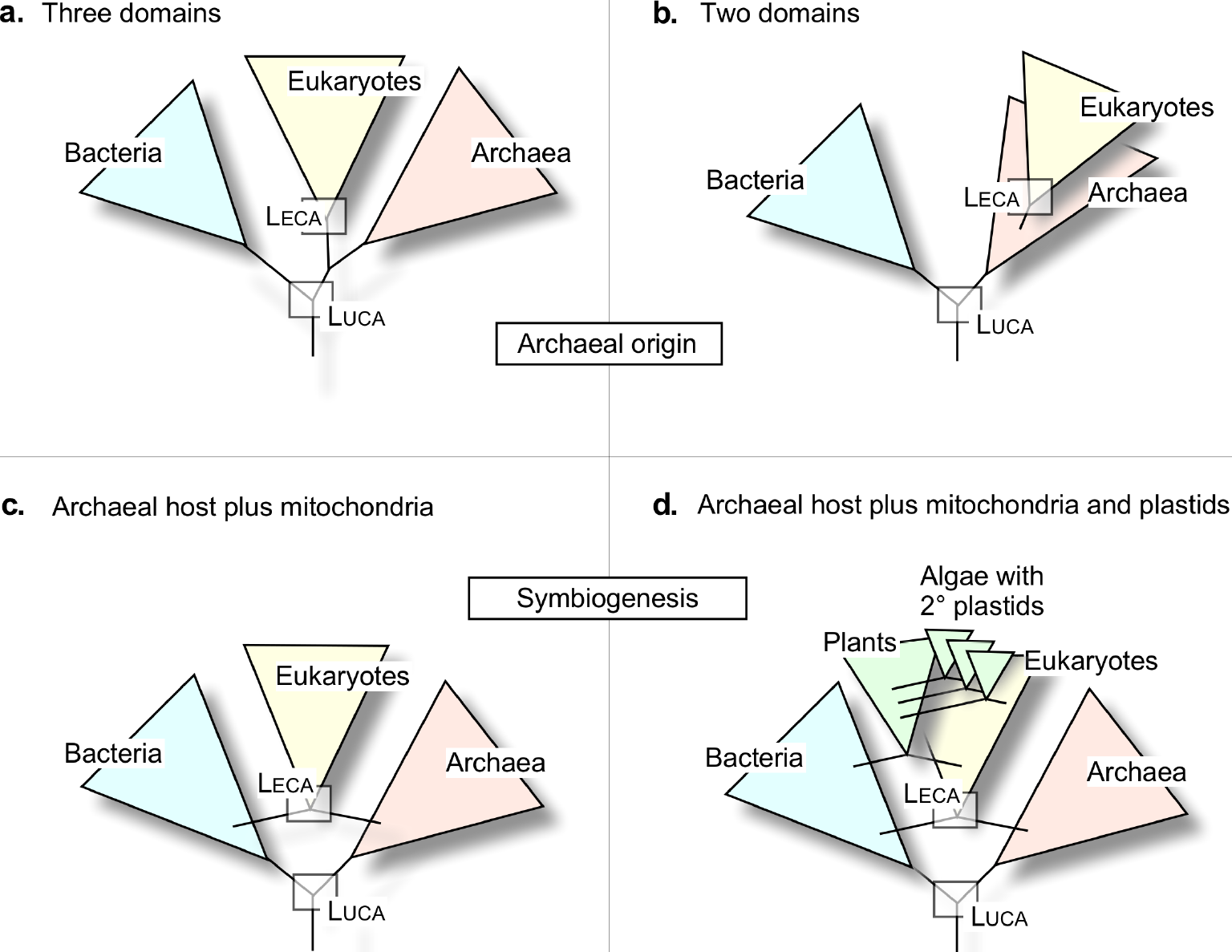
Differing views on the relationships of eukaryotes to prokaryotes. a) The three domain tree. b) The two domain tree with an archaeal origin of eukaryotes. c) Symbiogenesis at the origin of eukaryotes. d) Symbiogenesis at the origin of eukaryotes plus plastds at the origin of the plant kingdom and secondary symbiotic events among algae (see refs. [6, 23, 28, 34]).

Even though it is uncontested that symbiotic mergers lie at the root of modern eukaryotic groups via the single origin of mitochondria, plants via the single origin of plastids, and at least three groups of algae with complex plastids via secondary symbiosis [29], anastomosing structures such as those depicted in Fig. 1c and Fig. 1d do not mesh well with established principles of phylogenetic classification, because the classification of groups that arise by symbiosis is not unique. One could rightly argue that plants are descended from cyanobacteria, which is in part true because many genes in plants were acquired from the cyanobacterial antecedent of plastids [30]. Or one could save phylogenetic classification of eukaryotes from symbiogenic corruption by a democratic argument that eukaryotes are, by majority, archaeal based on the assumption that their genomes contain a majority of archaeal genes, making them archaea in the classificatory sense.

But what if eukaryotes are actually bacteria in terms of their genomic majority? The trees that molecular phylogeneticists use to classify eukaryotes are based on rRNA or proteins associated with ribosomes — cytosolic ribosomes in the case of eukaryotes. Ribosomes make up about 40% of a prokaryotic cell’s substance by dry weight, so they certainly are important for the object of classification. No one would doubt that eukaryotes have archaeal ribosomes in their cytosol. Archaeal ribosomes in the cytosol could however equally be the result of a gradualist origin of eukaryotes from archaea [31,32] or symbiogenesis involving an archaeal host for the origin of mitochondria [33,34]. Ribosomes only comprise about 50 proteins and three RNAs, while the proteins used for phylogenetic classification are only about 30 in number, or roughly 1% of an average prokaryotic genome [35]. The other 99% of the genome are more difficult to analyze, bringing us back to the question: At the level of whole genomes, are eukaryotes fundamentally archaeal?

Since the availability of complete genome sequences there have been investigations to determine the proportion of archaeal and bacterial genes in eukaryotic genomes. Such an undertaking is straightforward for an individual eukaryotic genome, and previous investigations have focused on yeast [36,37]. These indicated that yeast harbors an excess of bacterial genes relative to archaeal genes. Subsequent genome-wide phylogenetic analyses including plants, animals, and fungi [38,39], two eukaryotic groups [40] or six eukaryotic supergroups [41] fell short of reporting estimates for the proportion of genes in eukaryotic genomes that stem from bacteria and archaea respectively, whereby all previous estimates have been limited by the small archaeal sample of sequenced genomes for comparison. Here we have clustered genes from sequenced genomes of 150 eukaryotes, 5,443 bacteria and 212 archaea. By normalizing for the large bacterial sample through downsampling, we obtain estimates for the proportion of genes in each eukaryote genome that identify prokaryotic homologues, but that only occur in archaea or bacteria respectively.

## Results and Discussion

Using the MCL algorithm, we generated clusters for 19,050,992 protein sequences from 5,443 bacteria and 212 archaea with 3,420,731 protein sequences from 150 eukaryotes (see Methods) (Suppl. Table 1a-c) spanning six eukaryotic supergroups (Fig. 2a). This yielded 2,587 eukaryote-prokaryote clusters (EPC), 1,853 of which contained only eukaryotes and bacteria, 515 of which contained only eukaryotes and archaea. Among the 2,587 EPC clusters, 8% (219) contained sequences from at least two eukaryotes and at least five prokaryotes spanning bacteria and archaea (see Suppl. Table 2), which were not considered further for our estimates because here we sought estimates where the decision regarding bacterial or archaeal origin was independent of phylogenetic inference, which is possible for 92% of eukaryotic clusters that contain prokaryotic sequences. All sequences had unique cluster assignments, no sequences occurred in more than one cluster. That 1,853 clusters contained only eukaryotes and bacteria while 515 contained only eukaryotes and archaea appears to suggest a 3.6-fold excess of bacterial genes in eukaryotes, but bacterial genes are 25-fold more abundant in the data. For those genes that each eukaryote shares with prokaryotes, we estimated the proportion and number of genes having homologues only in archaea and only in bacteria respectively by downsampling the 25-fold excess of bacterial genomes in the sample in 1,000 subsamples of 212 bacteria and 212 archaea.

**Figure 2.**
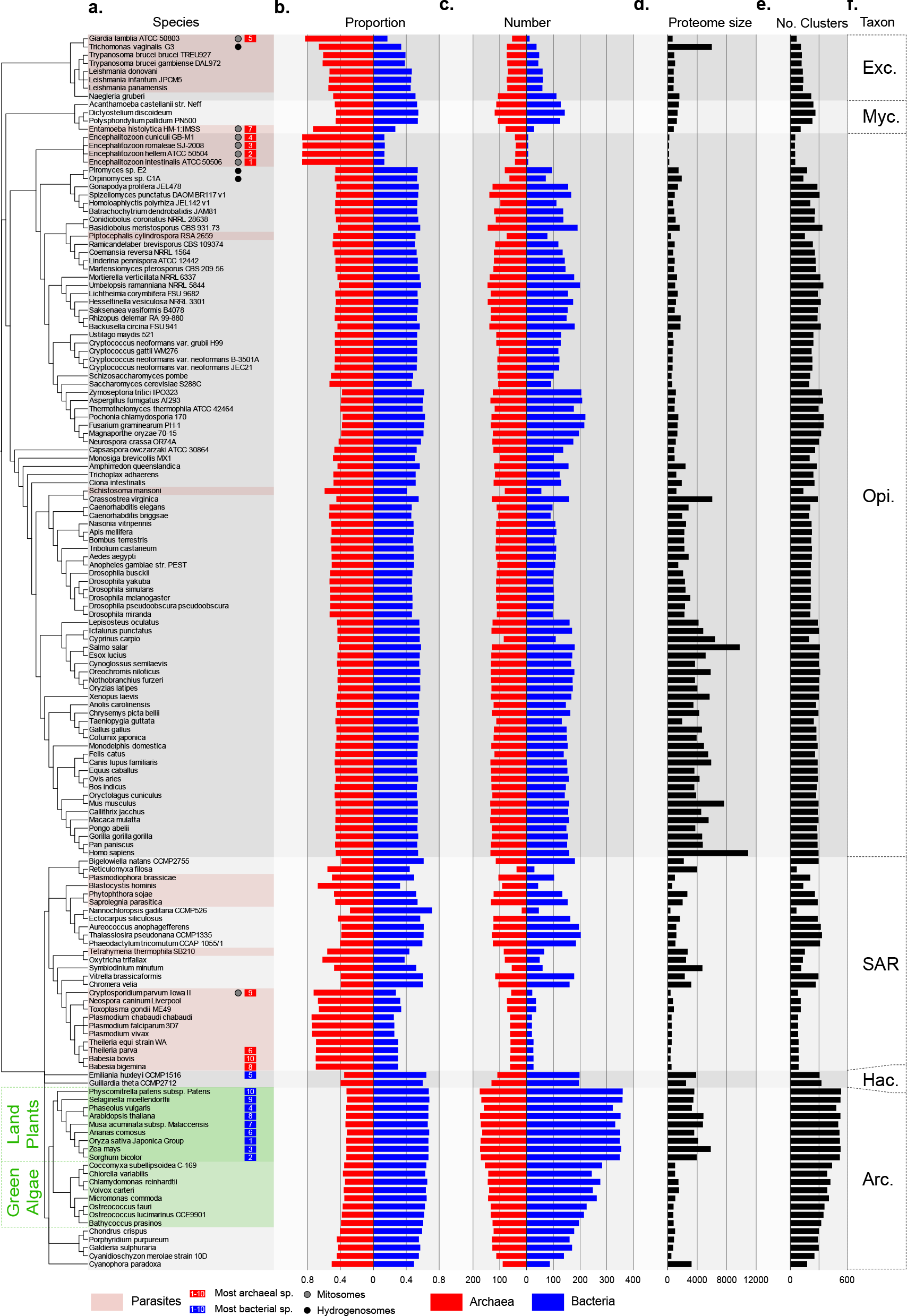
Bacterial and archaeal genes in eukaryotic genomes. Protein sequences from 150 eukaryotic genomes and 5,655 prokaryotic genomes (5,433 bacteria and 212 archaea) were clustered into eukaryote-prokaryote clusters (EPC) using the MCL algorithm [54] as described [41]. To account for overrepresentation of bacterial sequences in the clusters, bacterial genomes were downsampled in 1,000 datasets of 212 randomly selected bacterial organisms, the means were plotted. The eukaryotic sequences in the EPCs that cluster exclusively with bacterial or archaeal homologues were labelled bacterial (blue) or archaeal (red) accordingly. a.) Eukaryotic lineages and genomes were grouped by taxonomy. Numbers next to the species name on the left side indicate the ten most bacterial (blue) and archaeal (red) genomes, respectively. b.) The avg. relative proportion of bacterial and archaeal genes per genome. c.) The number of eukaryotic clusters with bacterial or archaeal homologs is shown. d.) The proteome size for the genome. e.) The sum of all eukaryotic sequences in the eukaryote-prokaryote clusters. Taxonomic groups are labelled on the far right panel. (Arc. – Archaeplastida, Exc. – Excavata, Hac. – Hacrobia, Myc. – Mycetozoa, Opi. – Opisthokonts). Highlighted in green is the branch with the taxons of plants and green algae, parasites are highlighted in red. The black dots indicate organisms with hydrogenosomes, the grey dot indicates organisms with mitosomes.

The proportion of bacterial and archaeal genes for each eukaryote is shown in Fig. 2b. Overall, 44% of eukaryotic sequences are archaeal in origin and 56% are bacterial. Across 150 genomes, eukaryotes possess 12% more bacterial genes than archaeal genes. There are evident group specific differences (Fig. 2b). If we look only at organisms that never harbored a plastid, the excess of bacteria genes drops from 56% to 53%. If we look only at groups that possess plastids the proportions of bacterial homologues increases to 61% vs. 39% archaeal (Table 1, Suppl. Table 3). Note that our estimates are based on the number of clusters, meaning that gene duplications do not figure into the estimates. A bacterial derived gene that was amplified by duplication to 100 copies in each land plant genome is counted as one bacterial derived gene. This is seen in Fig. 2 for Trichomonas, where a large number on gene families have expanded in the Trichomonas lineage [42], reflected in a conspicuously large proteome size (Fig. 2d), but a similar number of clusters (Fig. 2e) as neighboring taxa.

**Table 1.** Proportion of bacterial and archaeal derived genes in eukaryotic genomes.

| Group | Archaeal | Bacterial |
| --- | --- | --- |
| All eukaryotes | 0.44 | 0.56 |
| All without plastids<sup>1</sup> | 0.47 | 0.53 |
| All with plastids<sup>2</sup> | 0.39 | 0.61 |
| Land Plants | 0.33 | 0.67 |
| Opisthokonts | 0.46 | 0.54 |
| Hacrobia | 0.38 | 0.62 |
| SAR | 0.50 | 0.50 |
| Archaeplastida | 0.36 | 0.64 |
| Mycetozoa | 0.50 | 0.50 |
| Excavata | 0.58 | 0.42 |
| Parasites<sup>3</sup> | 0.62 | 0.38 |

Notes: <sup>1</sup> All except members of SAR, Hacrobia, and Archaeplastida as designated in supplementary table 3. <sup>2</sup> All members of SAR, Archaeplastida, and Hacrobia as designated in supplementary table 3. <sup>3</sup> Eukaryotes scored as parasites are designated in Fig. 1.

The proportions for different eukaryotic groups is shown in Table 1. Land plants have the highest proportion of bacterial derived genes at 67%, or a 2:1 ratio of bacterial genes relative to archaeal. The eukaryote with the highest proportion of bacterial genes in our sample is rice, with 67.1% bacterial and 32.9% archaeal genes. The higher proportion of bacterial genes in plastid containing eukaryotes relative to other groups corresponds with the origin of the plastid and gene transfers to the nucleus [41]. The eukaryote with the highest proportion of archaeal genes in our sample are the human parasite Encephalitozoon intestinalis and the rabbit parasite Encephalitozoon cuniculi, with 86% archaeal and 14% bacterial derived genes. Parasitic eukaryotes have the largest proportions of archaeal genes, but not by novel acquisitions, rather by having lost large numbers of bacterial genes as a result of reductive evolution in adaptation to nutrient rich environments. This is evident in Fig. 2c, where the numbers of archaeal and bacterial genes per genome are shown. Parasites, with their reduced genomes, such as Giardia lamblia, Trichomonas vaginalis, or Encephalitozoon species appear more archaeal. The number of archaeal, or bacterial genes in an organism does not correlate with genome size (Suppl. Fig. 1, Pearson correlation coefficient: archaeal r^2^ = 0.38, bacterial r^2^ = 0.33).

Opisthokonts generally have a more even distribution of bacterial and archaeal homologs in their genomes but are still slightly more bacterial (54%, Table 1 and Suppl. Table 3). The black and grey dots in Fig. 2a indicate organisms that possess reduced forms of mitochondria, hydrogenosomes (black) or mitosomes (grey). The ten most archaeal or bacterial organisms are indicated by a red or blue rectangle, respectively. The most archaeal eukaryotes are all parasites (highlighted in red) and have undergone reductive evolution, also with respect to their mitochondria, which are often reduced to mitosomes (Fig. 2a). Nine of the ten most bacterial organisms in the sample are plants (highlighted in green) with the fifth most bacterial organism being one of the only two Hacrobia in the dataset.

The functional distinction that eukaryotic genes involved in the eukaryotic genetic apparatus and information processing tend to reflect an archaeal origin while genes involved in eukaryotic biochemical and metabolic processes tend to reflect bacterial origins [43,44] has been borne out for yeast [36,37] and small genome samples [39,40]. The distributions of eukaryotic genes per genome that have archaeal or bacterial homologs across the respective KEGG function category at the first level (metabolism, genetic information processing, environmental information processing, cellular processes, and organismal systems) are shown in Fig. 3. The category human diseases is not shown, as only very few proteins in the eukaryote-prokaryote clusters were so annotated. The categories genetic information processing (information) and metabolism account for 90% of all annotated eukaryotic sequences in the EPCs (Suppl. Table 4). In the category metabolism, 67.6% of eukaryotic genes are bacterial while 76.9% of EPCs involved in information are archaeal. The distinction between informational and metabolic genes first described for yeast appears to be valid across all eukaryotic genomes.

**Figure 3:**
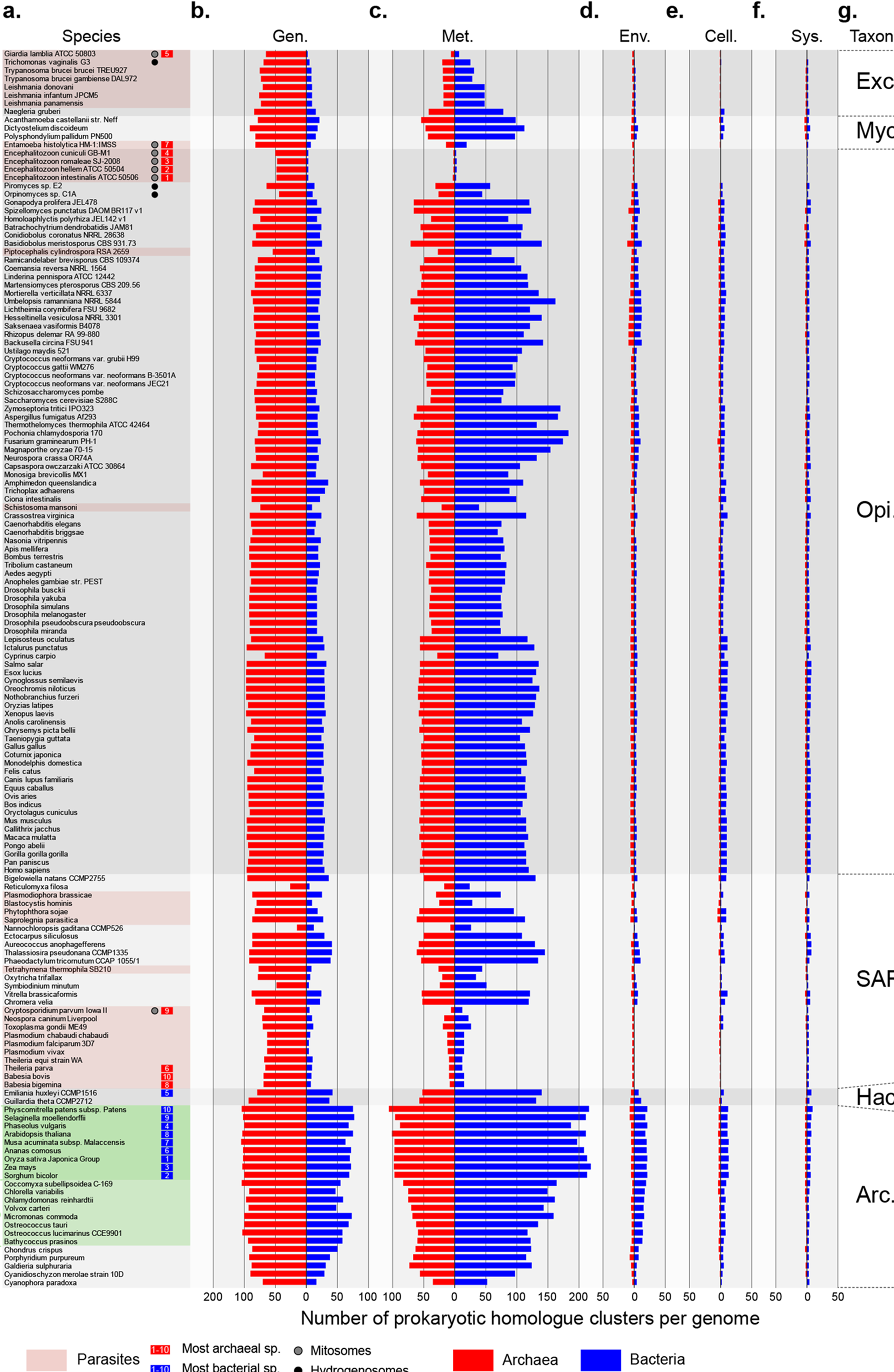
Functional categories. Protein sequences from 150 eukaryotic genomes and 5,655 prokaryotic genomes were clustered into 2,587 eukaryote-prokaryote clusters (EPC) [41]. Sorted according to a reference tree for eukaryotic lineages generated from the literature and taxonomic groups are labelled. The red bars indicate eukaryotic gene families that are archaeal in origin, blue indicates a bacterial origin of the gene family. Functional annotations according to the KEGG BRITE hierarchy on the level A was assigned for each EPC, identifying the function for each sequence in the protein cluster by performing a protein blast against the KEGG database and then applying the most prevalent function per protein family. Only the categories ‘Genetic Information Processing’ (Gen.), ‘Metabolism’ (Met.), ‘Environmental Information Processing’ (Env.), ‘Cellular Processes’ (Cell.), and ‘Organismal Systems’ (Sys.) are depicted, as the label ‘Human Diseases’ was hardly represented. Highlighted in green is the branch uniting land plants and green algae; the black and gray dots indicate organisms with hydrogenosomes or mitosomes, respectively.

The distribution of the genes in the 2,587 EPCs across genomes for six supergroups is depicted in Fig. 4. The order of eukaryotic and prokaryotic organisms (rows) can be found in Suppl. Table 5. Block A represents only Archaeplastida, block B depicts genes found in Archaeplastida and SAR, block C encompasses all genes that are distributed across the three taxa that contain plastids; Archaeplastida, SAR, and Hacrobia. The lower part of the figure shows the prokaryotic homologous genes. Cyanobacterial genes are especially densely distributed across blocks A–C. Genes that are predominantly mitochondrion- or host-related are indicated in block D and E. Eukaryotic genes that are universally distributed across the six supergroups are mainly archaeal in origin (block D). Especially organisms with reduced genomes such as parasites (marked with asterisks on the right), have lost genes associated with metabolism, leaving them mainly archaeal (Fig. 4). In the wake of symbiogenic mergers, which are very rare in evolution, gene loss sets in, whereby gene loss is very common in eukaryote genome evolution, one of its main underlying themes [41,45].

**Figure 4:**
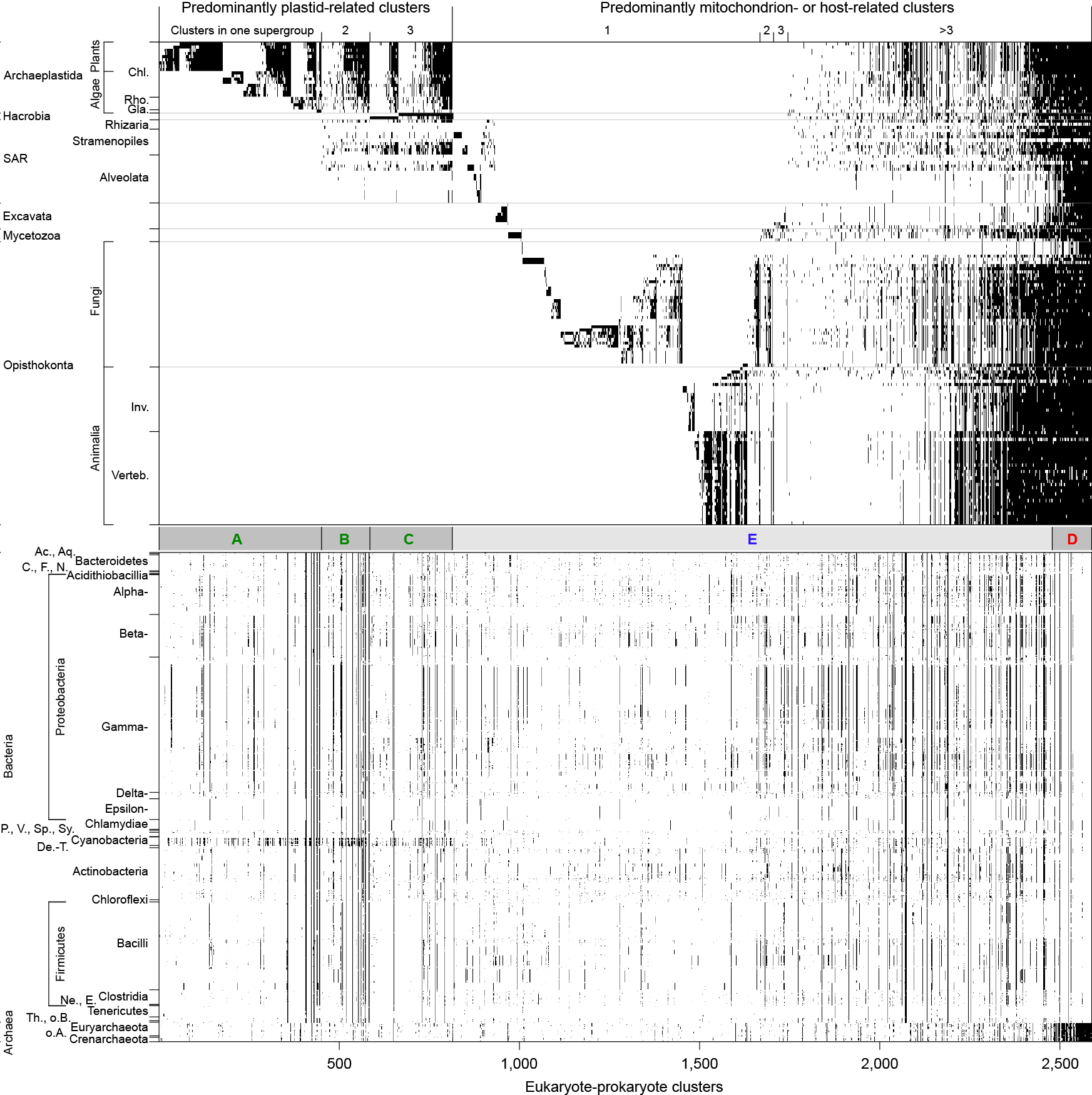
Gene sharing matrix. Each black tick represents the presence of a gene in the respective taxon. First, the 2,587 EPCs (x axis) were sorted according to their distribution across the six eukaryotic supergroups with the photosynthetic lineages on the left (block A–C). Host- or mitochondrion-related genes distributed across the six supergroups are depicted in block E. Clusters with mostly archaeal homologs are indicated in block D. (Chl. – Chloroplastida, Rho. – Rhodophyta, Gla. – Glaucophyta, Inv. – Invertebrates, Verteb. – Vertebrates; Ac. – Acidobacteria, Aq. – Aquificiae, C. – Chlorobi, F. – Fusobacteria, N. – Nitrospirae, P. – Planctomycetes, V. – Verrucomicrobia, Sp. – Spirochaetes, Sy. – Synergistetes, De.-T. – Deinococcus-Thermus, Ne. – Negativicutes, E. – Erysipelotrichia, Th. – Thermotogae, o.B. – other Bacteria, o.A. – other Archaea).

The estimates we obtain are based on a sample of genes that meet the clustering thresholds employed here. Many eukaryotic genes are inventions of the eukaryotic lineage in terms of domain structure and sequence identity. Those genes either arose in eukaryotes de novo from noncoding DNA, or they arose through sequence divergence, recombination, and duplication involving preexisting coding sequences, the bacterial and archaeal components of which should reflect that demonstrable in the conserved fraction of genes analyzed here. It is possible that archaeal genes and domains are more prone to recombination and rapid sequence divergence than bacterial domains are, but the converse could also be true and there is no a priori evidence to indicate that either assumption applies across eukaryotic supergroups. Hence with some caution, our estimates, which are based on the conserved fraction of sequences only, should in principle apply for the archaeal and bacterial components of the genome as a whole.

## Conclusion

Using a sample of 5,655 prokaryotic and 150 eukaryotic genomes and downsampling procedures to correct for the overabundance of bacterial genomes versus archaeal genomes for comparisons, we have obtained estimates for the proportion of archaeal and bacterial genes per genome in eukaryotes based on gene distributions. We found that the members of six eukaryotic supergroups possess a majority of bacterial genes over archaeal genes. If eukaryotes were to be classified by genome based democratic principle, they would be have to be grouped with bacteria, not archaea. The excess of bacterial genes disappears in the genomes of intracellular parasites with highly reduced genomes, because the bacterial genes in eukaryotes underpin metabolic functions that can be replaced by metabolites present in the nutrient rich cytosol of the eukaryotic cells that parasites infect. The functions of the ribosome and genetic information processing cannot be replaced by nutrients, hence reductive genome evolution in parasites leads to preferential loss of bacterial genes and leaves archaeal genes remaining. In photosynthetic eukaryote lineages, the genetic contribution of plastids to the collection of nuclear genomes is evident in our analyses, both in lineages with primary plastids descended directly from cyanobacteria and in lineages with plastids of secondary symbiotic origin. The available sample of archaeal genomes is still limiting for comparisons of the kind presented here.

As improved culturing and sequencing of complete archaeal genomes progresses, new lineages are being characterized at the level of scanning electron microscopy that branch, in ribosomal trees, as sisters to the host lineage at eukaryote origin [21]. These archaea are however not complex like eukaryotes, rather they are prokaryotic in size and shape and unmistakably prokaryotic in organization [21]. That is, the closer microbiologists hone in on the host lineage for the origin of mitochondria, the steeper the evolutionary grade between prokaryotes and eukaryotes becomes, in agreement with the predictions of symbiotic theory [21] (Fig. 5) and in contrast to the expectations of gradualist theories for eukaryote origin [33]. At the same time, the analyses presented here uncover a bacterial majority of genes in eukaryotic genomes, a majority that traces to the eukaryote common ancestor [41], which is also in line with the predictions of symbiotic theory. The most likely biological source of the bacterial majority of genes in the eukaryote common ancestor is the mitochondrial endosymbiont [41]. Genomes record their own history. Eukaryotic genomes testify to the role of endosymbiosis in evolution.

**Figure 5:**
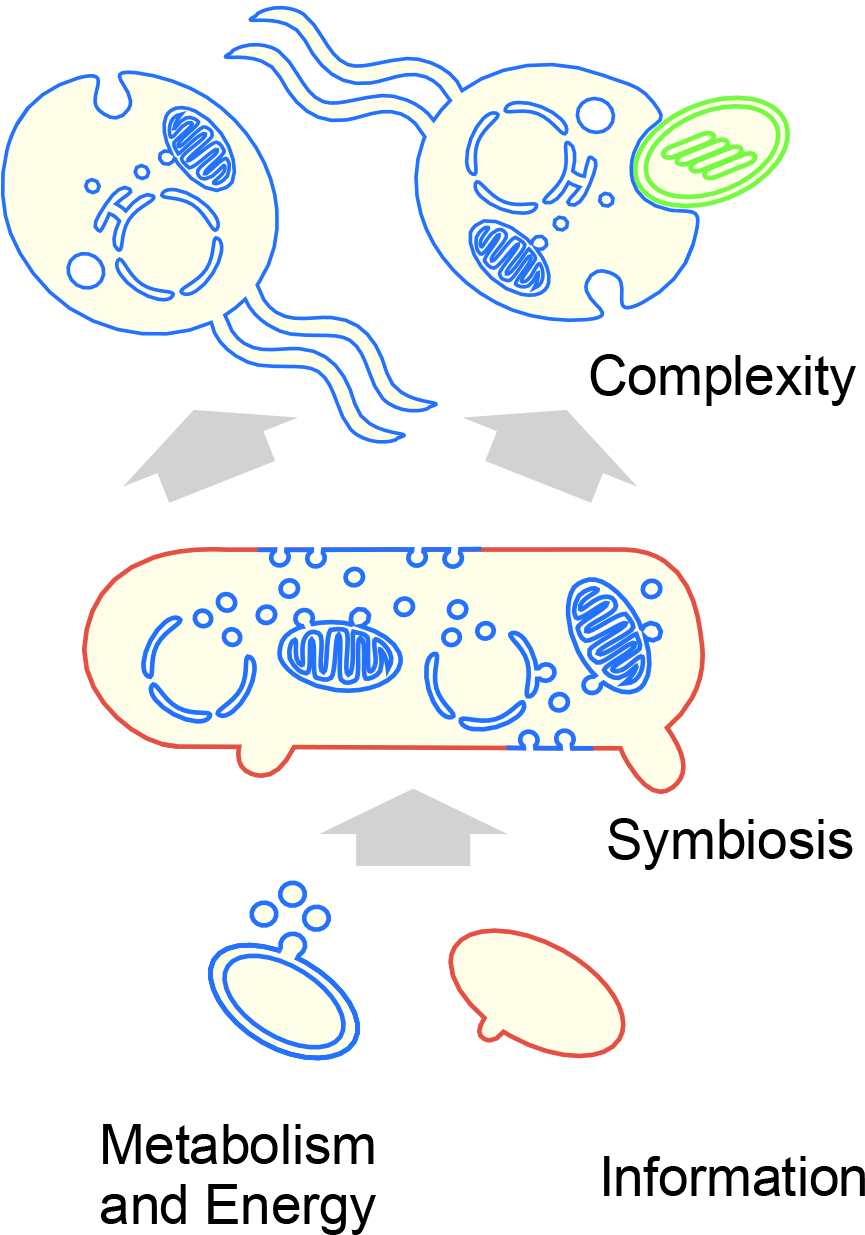
Bacterial and archaeal contributions to eukaryotes. Schematic representation of eukaryote origin involving an archaeal host and a mitochondrial symbiont that transforms the host via gene transfer from the endosymbiont [21,43]. The model combines elements of different proposals: bacterial outer membrane vesicles at the origin of the eukaryotic endomembrane system [4]; archaeal outer membrane vesicles at the origin of host membrane protrusions enabling endosymbiosis without phagocytosis [21]; a syncytial eukaryote common ancestor [56]; eukaryote origin starting an archaeal host and a bacterial symbiont brought into physical symbiotic interaction by anaerobic syntrophic interactions [21,43]; a combination of information (host) plus metabolism and energy (symbiont) [26,34] at eukaryote origin.

## Methods

### Sequence clustering

A total of 19,050,992 protein sequences from 5,655 complete prokaryotic genomes were downloaded from the NCBI RefSeq genomes database Release 78, September 2016 [46], encompassing 5,443 bacteria and 212 archaea (Suppl. Table 1a, 1b). For eukaryotes 3,420,731 protein sequences from 150 sequenced genomes covering a phylogenetically diverse sample were downloaded from NCBI RefSeq [46], Ensembl Protists [47], JGI [48], and GenBank [49] (Suppl. Table 1a, 1c) as appropriate. Protein sequences from the three domains were each clustered separately and homologous clusters were combined as described previously [41,50]. The reciprocal best BLAST hits (rBBH) [51] of an all-vs-all BLAST (v. 2.5.0) [52] were calculated for each domain (cut-off: expectation (E) value ≤ 1e−10). Pairwise global sequence identities were then generated for each sequence pair with the Needleman-Wunsch algorithm using the program ‘needle’ of the EMBOSS package v. 6.6.0.0 [53] with a global identity cut-off ≥ 25% for bacterial and archaeal sequence pairs and ≥ 40% global identity for eukaryotic sequence pairs. Protein families were reconstructed applying the domain-specific rBBH to the Markov Chain clustering algorithm (MCL) v. 12-068 [54] on the basis of the global pairwise sequence identities, respectively. Due to the large bacterial dataset, pruning parameters of MCL were adjusted until no relevant split/join distance between consecutive clusterings was calculated by the ‘clm dist’ application of the MCL program family (-P 180,000 −S 19,800 −R 25,200). MCL default settings were applied for the archaeal and eukaryotic protein clustering. This yielded 16,875 archaeal protein families (422,054 sequences) and 214,519 bacterial protein families (17,384,437 sequences) with at least five sequences each and 239,813 eukaryotic protein families (1,545,316 sequences) with sequences present in at least two species (Suppl. Table 6). To combine eukaryotic clusters with bacterial or archaeal clusters, the reciprocal best cluster approach [41] was applied with 50% best-hit correspondence and 30% BLAST local pairwise sequence identity of the inter-domain hits between eukaryote and prokaryote sequences. Eukaryotic clusters having homologues in both bacterial and archaeal clusters were merged with their prokaryotic homologues as described [41]. The cluster merging procedure left 752 eukaryotic clusters that had ambiguous (multiple) prokaryote cluster assignment, these were excluded from further analysis and 236,474 eukaryote clusters connected to no homologous prokaryotic cluster (eukaryote-specific, ESC, Suppl. Table 2) at the cut-offs employed here.

### Assignment of bacterial or archaeal origin

Because the number of prokaryotic sequences clustered was large, the 2,368 EPCs that were assigned one bacterial or one archaeal cluster exclusively were rechecked for homologs from the remaining prokaryotic domain at the E value ≤ 1e−10, global identity ≥ 25% threshold. The 266 cases so detected were excluded from bacterial-archaeal origin assignment, yielding 2,102 EPCs (Suppl. Table 2, indicated by asterisks). The clusters generated from rBBH (E value ≤ 1e−10, global identity ≥ 25%) of all-vs-all BLAST of the 19,050,992 prokaryotic protein sequences are provided as supplementary material (Suppl. Table 6). Downsampling to adjust for the overrepresentation of bacterial strains in the prokaryotic dataset compared to the number of archaeal organisms was performed by generating 1,000 datasets with 212 bacterial taxa selected randomly according to the distribution of genera in the whole dataset (Suppl. Table 7). The sequences of the examined 212 archaeal and bacterial taxa were located in the 2,102 EPCs and each eukaryotic organism in the identified clusters was assigned to ‘bacterial’, or ‘archaeal’ depending on the domain of the prokaryotic cluster in the EPC. Each eukaryotic genome was only counted once per EPC and assigned the respective prokaryotic label to prevent overrepresentation of duplication rich organisms. This procedure was performed for all 1,000 downsized bacterial datasets for each EPC, the mean of 1,000 samples was scored (Suppl. Table 3).

### Cluster annotation

Protein annotation information according to the BRITE (Biomolecular Reaction pathways for Information Transfer and Expression) hierarchy was downloaded from the Kyoto Encyclopedia of Genes and Genomes (KEGG v. September 2017) website [55], including protein sequences and their assigned function according to the KO numbers (Suppl. Table 8). The sequences of each protein family from the 2,587 EPCs were locally aligned with ‘blastp’ to the KEGG database to identify the annotation for each protein. In order to assign each protein to a KEGG function, only the best blast hit of the given protein with an E value ≤ 1e−10 and alignment coverage of 80% was selected. After assigning a function based on the KO numbers of KEGG for each protein in the EPCs, the majority rule was applied to identify the function for each cluster. The occurrence of the function of each protein was added and the most prevalent function was assigned for each cluster (Suppl. Table 4). Poorly characterized sequences or sequences with no assigned function were ignored, resulting in 1,836 clusters with annotations.

### Presence and absence of EPCs across genomes

Presence of absence of genes in a cluster for each genome were plotted as a 2,587 × 5,805 binary matrix, rows were sorted taxonomically, columns were sorted in ascending order left to right according to density of distribution within eukaryotic groups. Hacrobia and SAR were treated as a eukaryotic group for clusters they shared with Archaeplastida only; these clusters reflect secondary symbioses [41].

## Acknowledgments

We thank the European Research Council (grant 666053), and the Volkswagen Foundation (grant 93 046) for financial support. We thank Nils Kapust, Michael Knopp, Sriram Garg, Josip Skejo, Verena Zimorski and Sven Gould for helpful discussions.

